# *In vivo* calcium imaging visualizes peripheral neuron sensitization in murine osteoarthritis

**DOI:** 10.1101/160416

**Authors:** Rachel E. Miller, Yu Shin Kim, Phuong B. Tran, Xinzhong Dong, Richard J. Miller, Anne-Marie Malfait

**Author notes:** Corresponding authors: Anne-Marie Malfait, MD, PhD; Rachel E. Miller, PhD.

## Abstract

**Objective:** The purpose of this study was to develop a method for analyzing sensory neuron responses to mechanical stimuli *in vivo*, and to evaluate whether these neuronal responses change after destabilization of the medial meniscus (DMM).

**Methods:** DMM or sham surgery was performed in 10-week old male C57BL/6 wild-type or Pirt-GCaMP3^+/−^ mice. All experiments were performed eight weeks after surgery. Knee and hind paw hyperalgesia were assessed in wild-type mice. The retrograde label DiI was injected into the ipsilateral knee to quantify the number of knee-innervating neurons in the L4 dorsal root ganglion (DRG) in wild-type mice. *In vivo* calcium imaging was performed on the ipsilateral L4 DRG of Pirt-GCaMP3^+/−^ mice as mechanical stimuli (paw pinch, knee pinch, knee twist) were applied to the ipsilateral hind limb.

**Results:** Eight weeks after surgery, DMM mice had more hyperalgesia in the knee and hind paw compared to sham mice. Intra-articular injection of DiI labeled similar numbers of neurons in the L4 DRG of sham and DMM mice. Increased numbers of sensory neurons responded to all three mechanical stimuli in DMM mice, as assessed by *in vivo* calcium imaging. The majority of responses in sham and DMM mice were in small-to-medium-sized neurons, consistent with the size of nociceptors. The magnitude of responses was similar between sham and DMM mice.

**Conclusions:** We demonstrated that increased numbers of small-to-medium sized DRG neurons respond to mechanical stimuli 8 weeks after DMM surgery, suggesting that nociceptors have become sensitized by lowering the response threshold.

## Introduction

Nervous system sensitization, as determined by quantitative sensory testing, is associated with osteoarthritis (OA) and has been shown to correlate with symptom severity (1, 2). In knee OA, sensitization is associated with the presence of inflammation (3). A number of studies in small cohorts have suggested that following hip or knee joint replacement, sensitization is often reversed and this is associated with symptom relief (4-6).

Sensitization is also a feature of experimental OA, and can be detected by evaluating pain-related behaviors in animals, including mechanical allodynia and mechanical hyperalgesia of the hind limb (for review (7)). Previous work has shown that after destabilization of the medial meniscus (DMM), mice develop slowly progressive joint damage concurrent with pain-related behaviors indicative of sensitization, including hind paw mechanical allodynia and knee hyperalgesia, which develop prior to the manifestation of spontaneous pain behaviors such as locomotive deficits (8-11). In order to develop targeted analgesic strategies, the neuronal mechanisms that mediate this sensitization to mechanical stimuli need to be defined. Therefore, we sought to develop a method for analyzing peripheral sensory neuron activity in response to mechanical stimuli in real-time *in vivo*.

*In vivo* electrophysiology enables monitoring responses in knee-innervating afferent nerve fibers while physical stimuli are applied to the knee, but only one fiber may be measured at a time (12). In contrast, recently developed *in vivo* calcium imaging methods allow the simultaneous visualization of calcium (Ca)_i_ responses of hundreds of neurons within the dorsal root ganglion (DRG) (13, 14). In the current study, we adapted this novel method in order to visualize sensitization of knee and hind paw afferents after DMM or sham surgery in response to three types of physical stimuli applied to the hind limb. Operated mice were subjected to a low-level non-noxious stimulus below the threshold that would normally elicit a behavioral response, first through a paw pinch; and secondly, through a knee pinch. We also applied a noxious stimulus above the threshold that would elicit a behavioral response, by applying a noxious knee twist. We monitored neuronal responses in the L4 DRG by real-time *in vivo* calcium imaging. We focused on the 8-week time point after surgery because we have previously shown this to be a transition period from a state of acute to chronic pain (10, 15, 16), and it is when all pain-related behaviors associated with mechanical stimuli are apparent in the model (10, 11, 15).

## Methods

### Animals

A total of 85 mice were used. All animal experiments were approved by the Institutional Animal Care and Use Committees at Rush University Medical Center, Northwestern University, and Johns Hopkins University. Animals were housed with food and water *ad libitum* and kept on 12-hour light cycles. C57BL/6 wild-type and Pirt-GCaMP3^+/−^ mice were bred at Rush. Pirt-GCaMP3 mice express the fluorescent calcium indicator, GCaMP3, in ∼90% of all sensory DRG neurons, and not in other peripheral or central tissues, through the Pirt promoter (13, 17).

### Surgery

DMM surgery was performed in the right knee of 10-week old male mice (25 – 30 g), as previously described (10, 18), under isoflurane anesthesia. Briefly, after medial parapatellar arthrotomy, the anterior fat pad was dissected to expose the anterior medial meniscotibial ligament, which was severed. The knee was flushed with saline and the incision closed. Sham surgery followed the same procedure to expose the anterior medial meniscotibial ligament, but the ligament was left intact. Mice were not administered analgesia after surgery.

### Knee and Hind Paw Hyperalgesia

Knee hyperalgesia was measured in wild-type mice (n=7 naïve; n=6 sham; n=9 DMM) using a Pressure Application Measurement (PAM) device (Ugo Basile, Varese, Italy) as previously described (15, 19, 20). Briefly, mice were restrained by hand and the hind paw was lightly pinned with a finger in order to hold the knee in flexion at a similar angle for each mouse. With the knee in flexion, the PAM transducer was pressed against the medial side of the ipsilateral knee while the operator’s thumb lightly held the lateral side of the knee. The PAM software guided the user to apply an increasing amount of force at a constant rate (30 g/s), up to a maximum of 450 g. If the mouse tried to withdraw its knee, the force at which this occurred was recorded. If the mouse did not try to withdraw, the maximum possible force of 450 g was assigned. Two measurements were taken per knee and the withdrawal force data were averaged. Hind paw hyperalgesia was measured in the same way (n=9 naïve; n=5 sham; n=8 DMM), but the PAM transducer was applied to the ventral aspect of the ipsilateral hind paw with the operator’s thumb against the dorsal aspect.

### In vivo calcium imaging

Six to seven weeks after surgery, Pirt-GCaMP3^+/−^ mice were shipped from Rush to Johns Hopkins University and were allowed to recover for one to two weeks. Eight weeks after surgery, mice were deeply anesthetized using sodium pentobarbital (40-50 mg/kg), and the back and operated knee were shaved. A laminectomy from vertebrae L2-L6 was performed, and the L4 DRG was exposed, as previously described (13). The L4 DRG was chosen since it contains the cell bodies of the majority of sensory neurons that innervate the knee of the mouse (21, 22). The mouse was positioned under a laser scanning confocal microscope (Leica LSI) by clamping the spinal column at L2 and L6 using forceps attached to micromanipulators. Anesthesia was maintained using isofluorane during imaging and temperature was maintained using a homeothermic blanket system. Images were acquired using a 0.5 N.A. macro dry objective and an EM-CCD camera (488 nm excitation, 500-550 nm emission). In order to capture the entire visible area of the DRG, z-stacks were taken at 600 Hz in 10 steps over a distance of 200-300 μm at 512x512 pixel resolution in the x-y plane. A time series was performed such that in total, approximately 15 z-stacks were taken for each stimulus. Each z-stack took approximately 7-8 seconds to achieve. For each mouse (n=12 sham, n=14 DMM), physical stimuli were applied to the ipsilateral limb in the following order: 1. A 100g force was applied to the hindpaw using a calibrated forceps system (IITC Rodent Pincher); 2. A 30g force was applied to the knee using the same calibrated forceps system; 3. A noxious outward rotation (∼60°, approximately 40-60 mNm torque (12), which is beyond the normal physiological range but does not cause joint injury (23, 24)) was applied to the knee by holding the femur in place with a forceps while the hind paw is used to rotate the lower leg (‘knee twistx2019;). One sham mouse was excluded from the knee twist analysis due to excessive movement during imaging. For each stimulus, baseline images were captured for 5-6 z-stacks (approximately 45 seconds) prior to the application of the stimulus, the stimulus was applied for 5 z-stacks, and an additional 4-5 z-stacks were captured after the stimulus was discontinued. Between each stimulus, the mouse was allowed to recover for at least 3 minutes in order to ensure that all previous neuron responses had ceased and the fluorescence levels had returned to baseline. Pilot experiments demonstrated that the order of the applied stimuli did not change the number of responses, indicating that these acute stimuli were not sufficient to induce sensitization and thus did not seem to affect the following stimulus applied. At the end of each experiment, a high-resolution z-stack was taken at 1024x1024 pixel resolution with 4-8 frame averaging to use for counting the total number of neurons.

### Analysis of *in vivo* calcium imaging

Using the Leica software, a maximum intensity projection was performed for each z-stack in a particular time series, and the files were exported for further processing in Fiji (25) using custom macros and the Multi Measure plug-in. Brightness and contrast were adjusted, and all videos were first analyzed by a blinded observer to identify responding cells by looking for an increase in fluorescence during the stimulus application period. All responding cells were labeled as a region of interest (ROI) for further analysis. Cells spontaneously responding prior to application of the stimulus were ignored – in general, there were few spontaneous responses and there was no difference between sham and DMM groups. In addition, responses that occurred upon release of the stimulus were ignored. The total number of neurons imaged for each DRG was estimated by counting the number of neurons within a region of average density and extrapolating to the total DRG area. In order to confirm the visual assessment, changes in (Ca)_i_ were quantified by calculating the change in fluorescence for each ROI in each frame t of a time series using the formula: ΔF/F_o_ = (F_t_ - F_o_)/F_o_, where F_o_ = the average intensity during the baseline period prior to the application of the stimulus. For each stimulus, the maximum ΔF/F_o_ and area under the curve was calculated for each responding cell in a particular DRG. In order to compare the sizes of responding cells between sham and DMM mice for a particular stimulus, the areas of the ROIs were calculated in Fiji. For each mouse, a frequency distribution using relative frequencies was computed for the responding cell areas using a bin range with bin centers from 150-1550 μm^2^ and a bin width of 100 μm^2^. Areas greater than 800 μm^2^ were summed into one bin. The calculated relative frequencies for each size category were averaged across the mice for either DMM or sham treatment.

### Histopathology of the knee

Following *in vivo* calcium imaging, mice were euthanized by carbon dioxide inhalation, and knees were collected for histopathology and were evaluated based on modified OARSI recommendations, as previously described (n=12 DMM; n=11 sham) (26) (Alison Bendele, Bolder BioPATH, Inc., Boulder CO). Joints were fixed in 10% formalin, decalcified, embedded in the frontal plane, sectioned (8 μm), and stained with Toluidine blue (0.04% w/v). A coronal section from the mid-joint (area of maximal damage (18)) was used to score the medial femoral condyles and tibial plateaux for severity of cartilage degeneration. For each cartilage surface, scores were assigned individually to each of 3 zones (inner, middle, outer) on a scale of 0-5, with 5 representing the most damage (maximal summed score for femoral + tibial cartilage degeneration = 30). The largest osteophyte (medial tibia or femur) was measured using an ocular micrometer.

### Retrograde Labeling and Immunofluorescence

Wild-type DMM mice 8 weeks post-surgery (n=6) along with age-matched naïve (n=3) and sham (n=6) controls were anesthetized with isoflurane and intra-articularly injected in the ipsilateral right knee with 5 μL DiI (2.5 mg/mL in methanol; ThermoFisher, D3911). One week post-injection, mice were anesthetized by ketamine and xylazine and perfused transcardially with PBS followed by 4% paraformaldehyde in PBS. The spinal column was dissected and postfixed in 4% paraformaldehyde overnight followed by cryopreservation in 30% sucrose in PBS. Ipsilateral L4 DRG were embedded with OCT (Tissue-Tek), frozen with dry ice, and cut into 12 μm sections. For immunostaining, slides were allowed to dry at room temperature for 2 h, postfixed with 4% paraformaldehyde for 10 min, and washed with PBS. Sections were blocked and permeabilized with 5% normal goat serum in 0.1% triton in PBS prior to incubation with PGP9.5 rabbit polyclonal antibody (Abcam ab27053; 1:200) overnight at 4 °C. Sections were washed with PBS and incubated with appropriately conjugated AlexaFluor 488 antibody (Invitrogen, 1:500) for 1 h at room temperature. Lastly, the sections were washed with PBS and mounted with Vectashield mounting media. DiI (549-565 nm excitation) and PGP9.5 signals were captured using a confocal microscope and the images were analyzed using ImageJ and Photoshop. For quantification, three sections per DRG were used. The number of neurons that expressed DiI were summed across the 3 sections and normalized to the total number of DRG neurons (PGP9.5+) summed across the 3 sections. The counts for DMM and sham mice were performed in a blinded fashion.

### Statistics

GraphPad Prism version 6.07 was used for statistical calculations. Data are expressed as either mean ± standard error of the mean (SEM) or median ± interquartile range (IQR), as indicated. For hyperalgesia experiments, groups were compared by one-way ANOVA followed by Tukey’s multiple comparisons test. A p-value < 0.05 was considered to be significant. For numbers of responding cells, clustering, nearest neighbor distance, change in fluorescence, and area under the curve, data were tested for normality by D’Agostino & Pearson omnibus normality test. If data passed the normality test, an unpaired t test was used; otherwise, a Mann-Whitney test was used. For size of responding cells to a particular stimulus, a two-way ANOVA followed by Sidak’s multiple comparisons test was used to compare average relative frequencies for each size category between sham and DMM mice, and a Mann-Whitney test was used to compare the median size of responding cells between sham and DMM mice. For retrograde labeling, groups were compared by Kruskal-Wallis test followed by Dunn’s multiple comparisons test.

## Results

### More neurons respond to mechanical stimuli applied to the hind limb 8 weeks after DMM than after sham surgery

Eight weeks after surgery, wild-type DMM mice had increased primary hyperalgesia in the knee joint compared to sham and naïve mice (Fig 1A), similar to our previous results (15), and this was accompanied by secondary hyperalgesia in the hind paw (Fig 1B). Both behaviors are indicative of sensitization. Therefore, we focused on this time point for all imaging experiments and imaged the L4 DRG, since these ganglia contain the majority of the cell bodies of sensory neurons that innervate the mouse hindlimb, including both the paw and the knee. For each DRG, we imaged a similar number of neurons for DMM (981±69) and sham (990±53) mice (p=0.9194) (Fig 2A – example images). Eight weeks after surgery, we observed that, for all 3 stimuli, an increased percentage of neurons in DMM mice responded compared to sham mice, as evidenced by an increase in transient (Ca)_i_ following stimulation (Fig 2B-D; Supplemental Videos 1-6). The noxious knee twist induced more responses than the 30-g knee pinch in sham mice (p=0.0199), but the two stimuli induced similar numbers of responses in DMM mice (p=0.1908). Following imaging, a subset of knees was analyzed to confirm joint damage. Similar to previous studies (10, 26), DMM mice developed moderate levels of cartilage damage (n=12; mean±SEM: 7.7±1.0) and osteophytes (159±16 μm) in the medial compartment by this time point, while sham mice did not develop joint damage (n=11; cartilage damage = 0±0); osteophyte width = 0±0).

**Figure 1:**
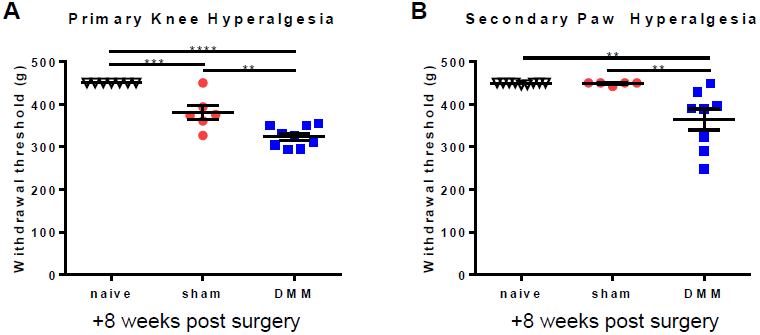
Wild-type mice that have undergone DMM surgery have increased (A) primary knee hyperalgesia and (B) secondary paw hyperalgesia 8 weeks after surgery compared to sham and age-matched naïve mice (**p<0.01, ***p<0.001, ****p<0.0001). mean±SEM.

**Figure 2:**
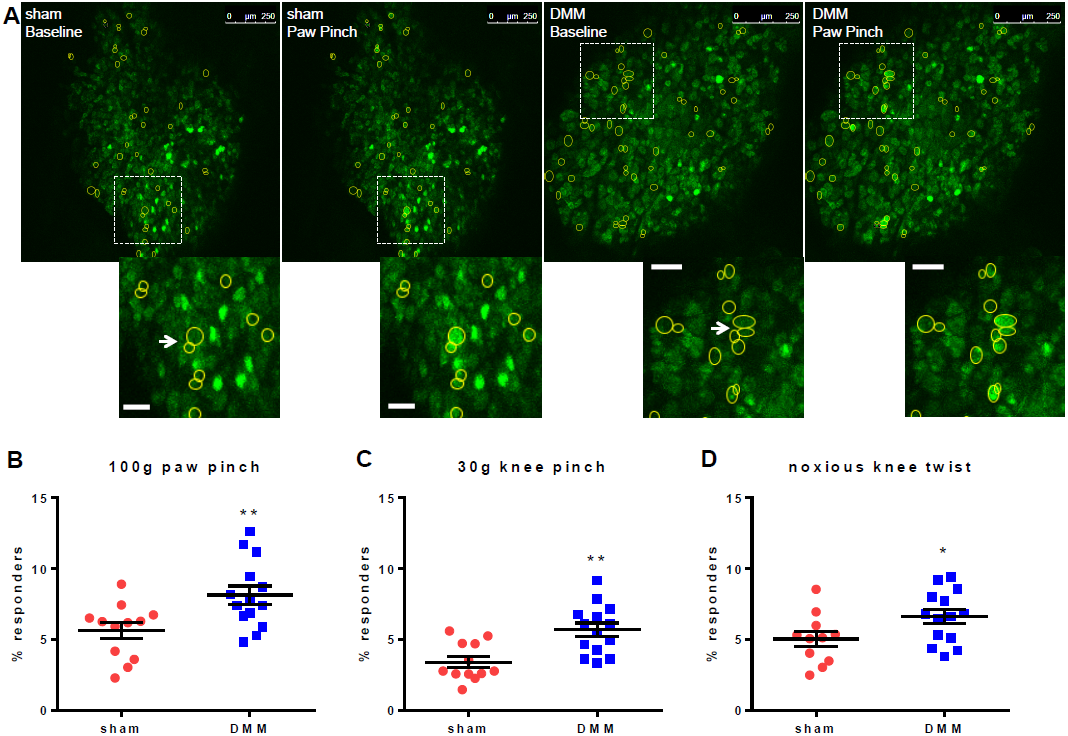
Eight weeks after surgery, paw- and knee-directed stimuli cause increased numbers of L4 DRG neuron responses in DMM mice compared to sham mice. (A) Representative images depicting L4 DRG neurons at baseline and responding to a 100 g pinch of the ipsilateral hind paw. Scale bars = 250 μm. Insets show examples of responding neurons shown with the arrow. The arrow also identifies examples of neuron clusters. Scale bar in insets = 50 μm. (B) Percent of neurons responding to 100 g ipsilateral paw pinch (# responding neurons / # total neurons in the field of view x 100). (C) Percent of neurons responding to 30 g ipsilateral knee pinch. (D) Percent of neurons responding to noxious knee twist. Each dot shows the percent of neurons that responded to the stimulus in one mouse L4 DRG. *p<0.05, **p<0.01. mean±SEM.

### Similar sized neurons respond in sham and DMM mice

In order to define which types of neurons responded before and after surgery, we assessed the size distribution of these cells. The majority of responses to all stimuli were in small-to-medium sized neurons (area < 600 μm^2^), consistent with the size of C and Aδ fiber neurons, the majority of which are nociceptors (27, 28) (relative frequencies shown in Fig 3A-C). However, there was no difference in the size distribution (Fig 3A-C) of responding neurons between sham and DMM mice for any of the three stimuli (p>0.8 for all comparisons). In addition, the median area was similar between sham and DMM mice for all three stimuli (paw pinch p=0.6770, knee pinch p=0.9798, knee twist p=0.8089) and was consistent with the size of small neurons (284-313 μm^2^) (Fig 3D). Together, these data suggest that C and Aδ fiber neurons have become sensitized by 8 weeks after DMM surgery, as opposed to larger Aβ fibers being recruited.

**Figure 3:**
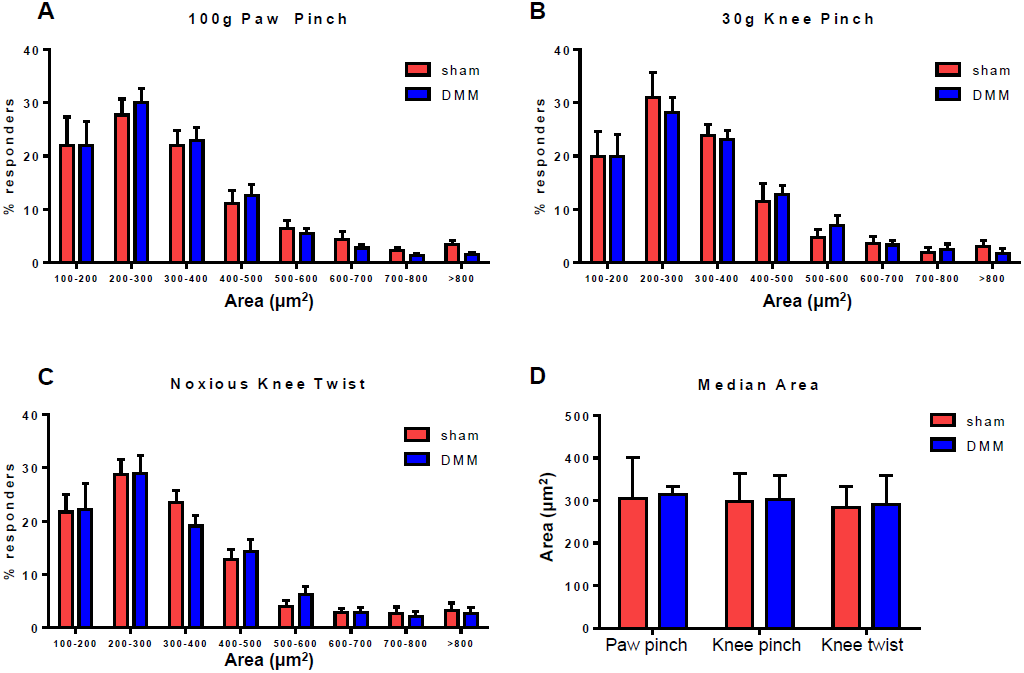
The size of responding neurons is similar in DMM and sham mice. (A-C) Relative frequency distributions of the areas of responding neurons in sham and DMM mice to (A) 100 g paw pinch (interaction of size and treatment (sham/DMM): p=0.9888), (B) 30 g knee pinch (interaction of size and treatment: p=0.9912), and (C) noxious knee twist (interaction of size and treatment: p=0.9474). mean±SEM. (D) Median area of responding cells. Sham *vs*. DMM, paw pinch: p=0.6770; knee pinch: p=0.9798; knee twist: p=0.8089. median±IQR.

### Similar numbers of neurons innervate the knee in sham and DMM mice

In order to exclude the possibility that the number of sensory neurons innervating the knee was increased 8 weeks after DMM, we performed retrograde labeling using DiI. Similar numbers of neurons were labeled in the L4 DRG of naïve mice 18 weeks of age and in sham and DMM mice 8 weeks after surgery (Table 1; Supp Fig 1). This result suggests that the number of sensory neurons innervating the knee remained the same over the course of the experiment.

**Table 1:** Percentage of L4 DRG Neurons Retrogradely-Labeled 7 Days after Intra-articular Injection of DiI.

| **Treatment** | **# labeled neurons** | **# total neurons** | **% labeled** | **median** | **p-value** |
| --- | --- | --- | --- | --- | --- |
| **Naïve** | 56 | 389 | 14.4 | 15.4 | >0.9999 vs sham and DMM |
|  | 79 | 445 | 17.8 |  |  |
|  | 78 | 506 | 15.4 |  |  |
| **Sham** | 29 | 478 | 6.1 | 14.3 | >0.9999 vs DMM |
|  | 48 | 565 | 8.5 |  |  |
|  | 48 | 423 | 11.3 |  |  |
|  | 77 | 403 | 19.1 |  |  |
|  | 62 | 358 | 17.3 |  |  |
|  | 78 | 448 | 17.4 |  |  |
| **DMM** | 99 | 627 | 15.8 | 15.3 |  |
|  | 60 | 415 | 14.5 |  |  |
|  | 98 | 648 | 15.1 |  |  |
|  | 64 | 409 | 15.6 |  |  |
|  | 108 | 399 | 27.1 |  |  |
|  | 65 | 462 | 14.1 |  |  |

### No change in the spatial organization of responses between sham and DMM mice

A recent report using this same *in vivo* calcium imaging technique demonstrated that in models of inflammatory and neuropathic pain, an increase in numbers of neuronal responses was associated with an increase in neuronal coupling (neurons immediately adjacent to each other responding to the same stimulus), which may represent a mechanism for neuronal sensitization through gap junctions (13). In contrast, in the current study, we found similar instances of neuronal coupling in both sham and DMM mice for all 3 stimuli (for an example of coupling, see arrows in insets of Fig 2A pointing to clusters of neurons). The percentage of coupled responding neurons (Fig 4A) as well as the mean number of neurons forming the clusters (Fig 4B) was similar between sham and DMM mice, suggesting that the formation of these clusters may be more associated with acute tissue injury as opposed to the chronicity of pain in this model. In addition, the median nearest neighbor distance of responding neurons was similar between sham and DMM mice for all stimuli (Fig 4C).

**Figure 4:**
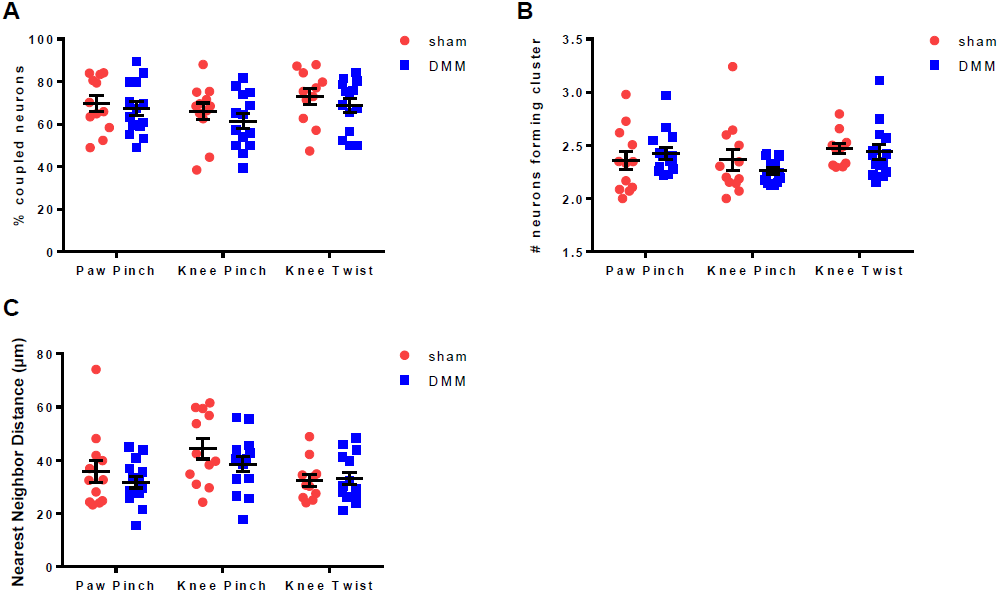
The spatial organization of responding neurons is similar in DMM and sham mice. (A) The percentage of responding neurons that were coupled in each DRG (# responding neurons touching at least one other responding neuron / total # responding neurons x 100). Sham *vs*. DMM, paw pinch: p=0.6222; knee pinch: p=0.3718; knee twist: p=0.3925. mean±SEM. (B) The mean number of responding neurons that made up an individual cluster in each DRG (# coupled responding neurons / # clusters x 100). Sham *vs*. DMM, paw pinch: p=0.4025; knee pinch: p=0.7136; knee twist: p=0.4030. median±IQR. (C) The mean nearest neighbor distance for responding neurons in each DRG. Sham *vs*. DMM, paw pinch: p=0.8596; knee pinch: p=0.4319; knee twist: p>0.9999. median±IQR.

### No change in the magnitude or duration of responding neurons

The intensity of each responding neuron was also assessed by quantifying the change in fluorescence with time (sample responses are shown in Fig 5A for responses to paw pinch in one DMM DRG). Overall, the maximum magnitude (Fig 5B) and duration (measured by area under the curve) (Fig 5C) of the responding population of neurons remained the same when comparing sham and DMM mice for all 3 stimuli, suggesting that the increased number of neurons recruited in the case of DMM mice is responsible for the increased sensitivity through lowering of the response threshold to these stimuli. For both sham and DMM mice, the noxious knee twist induced the largest magnitude and duration of response, which is consistent with the pain level induced by this stimulus (Fig 5B, C; Supp. Videos 5,6).

**Figure 5:**
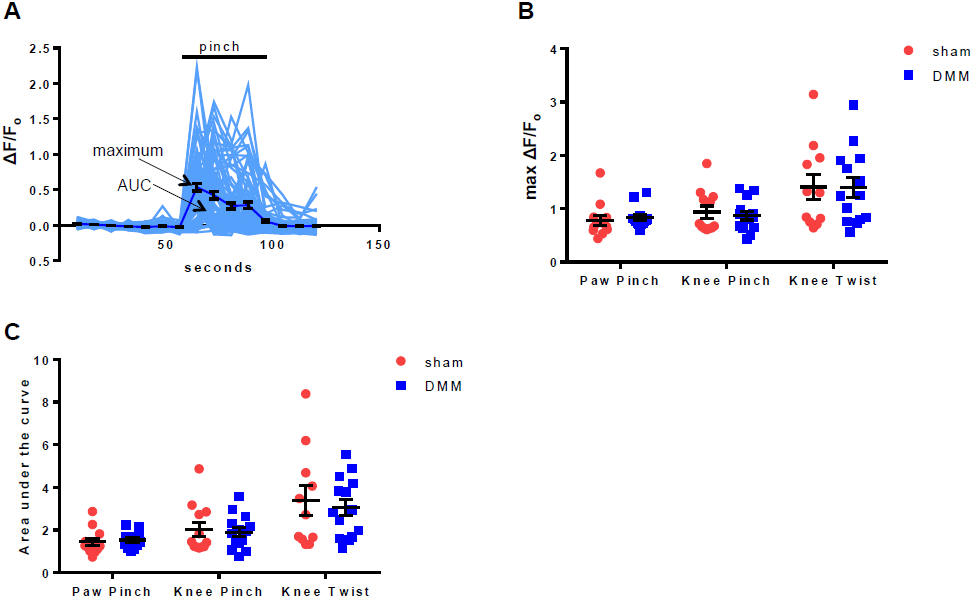
The magnitude and duration of (Ca)_i_ responses are similar in DMM and sham mice. (A) Individual traces (grey) and mean±SEM (black) of ΔF/F_o_ for neurons in a representative DMM DRG responding to 100 g paw pinch. (B) The mean maximum ΔF/F_o_ of responding neurons for each DRG. Sham *vs*. DMM, paw pinch: p=0.2312; knee pinch: p=0.6151; knee twist: p=0.9646. mean±SEM. (C) The mean area under the curve of responding neurons for each DRG. Sham *vs*. DMM, paw pinch: p=0.2740; knee pinch: p=0.8596; knee twist: p=0.6786. median±IQR.

## Discussion

We demonstrated that increased numbers of DRG neurons responded to physical stimuli directed toward the operated knee and ipsilateral hind paw in mice 8 weeks after DMM surgery compared to sham-operated mice, correlating well with pain-related behaviors. In addition, it was possible to analyze the size of the population of neurons that became sensitized in an unbiased fashion, since hundreds of neurons could be monitored simultaneously without determining *a priori* which type of fiber to record.

Overall, the majority of responses in both sham and DMM mice occurred in small-to-medium sized neurons, consistent with the size of the cell bodies of C and Aδ fibers. These fibers encompass both the nociceptor and C-low threshold mechanoreceptor populations (29). It is likely that increased (Ca)_i_ reflects increased excitability of these neuronal populations. Thus, it appears that at the 8-week time point after DMM surgery, C and Aδ fibers were sensitized such that increased numbers of neurons responded to subthreshold stimuli such as paw and knee pinches (forces below the threshold necessary to elicit a behavioral response), and also to a suprathreshold stimulus in the case of the knee twist (force that would normally cause a behavioral response). In addition, it does not appear that the increase in numbers of responses is due to recruitment of large numbers of large-diameter afferents (which are low threshold sensory neurons, not nociceptors (29)) such as the large-sized Aβ fibers, which have been implicated in mediating mechanical allodynia in nerve injury models (30) and have altered electrophysiology properties in a surgical rat model of OA (31). The relevance of these responses to OA pain is further highlighted by our recent observations that chemogenetic silencing of the Na_V_1.8 expressing population of DRG neurons, which primarily consist of C and Aδ fibers, inhibited mechanical allodynia at this same time point (8 weeks) after DMM surgery (15).

Our results are consistent with *in vivo* electrophysiology studies that have shown that noxious outward rotation of the knee joint causes increased firing of afferents in the knee joint compared to non-noxious rotation in healthy and monoiodoacetate (MIA) treated rats and guinea pigs (12, 24, 32). In addition, the fact that we observed increased responses in small-to-medium sized DRG neurons is consistent with electrophysiology studies on joint rotation in the MIA rat model of OA pain (32, 33). The firing rate of C and Aδ fiber afferents is elevated by day 14 after injection of MIA into knee joint compared to saline-injected controls, and this in response to both non-noxious and noxious rotation of the knee joint (32). In addition, the mechanical threshold of firing in response to rotation decreased in MIA rats (33).

While we observed a marked increase in the numbers of responding neurons after DMM surgery, we did not observe a change in the spatial organization of responding neurons compared to sham mice. A recent paper by Kim *et al*. reported that the amount of neuronal coupling seen in mice 2 days after injection of Complete Freund’s Adjuvant into the hind paw was elevated compared to the amount of coupling seen in naïve mice in response to a wide-range of hind paw pinch forces (13). This interneuronal coupling was mediated by communication through gap junctions, and the authors demonstrated that inhibition of connexin-43 expression in the DRG reduced the coupling as well as the pain behavior induced by this model. In the current study, we observed that the majority of neurons responding to paw pinch, knee pinch, or knee twist were adjacent to at least one other responding neuron, but we did not observe a difference in the extent of neuronal coupling between DMM and sham mice. Therefore, in this model, it does not appear that neuronal coupling in the DRG mediates the sensitization associated with the pain behaviors observed at this stage of the model, but rather may develop as a result of surgery. A limitation is that we were unable to directly compare to a non-surgical naïve group in the current study.

In conclusion, by adapting a novel *in vivo* imaging technique, we demonstrated that increased numbers of small-to-medium sized DRG neurons respond to mechanical stimuli 8 weeks after DMM surgery. This result suggests that nociceptor sensitization occurs through recruitment of additional nociceptors by decreasing the response threshold, which may contribute to the development of chronic pain behaviors in this model. It is clear therefore, that the use of methodology such as that introduced here will provide a powerful new way of screening novel therapeutic agents that treat OA pain by reducing the excitability of the set of nociceptors recruited in association with the disease.

## Acknowledgements

We would like to thank Dr. Kyoungsook Park, Dr. Qin Zheng, and Mr. Michael Anderson for assistance with imaging at Johns Hopkins University School of Medicine.

---

**Supplemental Figure 1:** Representative images of the L4 ipsilateral DRG of (A) naïve, (B) sham, and (C) DMM mice 7 days after retrograde label with DiI, which was injected into the ipsilateral knee joint intra-articularly. Neurons were counter-stained with PGP9.5, a pan-neuronal marker, for normalization purposes. Scale bar = 100 μm.

**Supplemental Video 1:** Representative video of a DMM DRG responding to paw pinch (2 fps).

**Supplemental Video 2:** Representative video of a sham DRG responding to paw pinch (2 fps).

**Supplemental Video 3:** Representative video of a DMM DRG responding to knee pinch (2 fps).

**Supplemental Video 4:** Representative video of a sham DRG responding to knee pinch (2 fps).

**Supplemental Video 5:** Representative video of a DMM DRG responding to knee twist (2 fps).

**Supplemental Video 6:** Representative video of a sham DRG responding to knee twist (2 fps).

